# IL-23 production is regulated via an MSK1/2 – CREB dependent signalling pathway downstream of Toll like receptors

**DOI:** 10.1101/2020.07.07.189142

**Authors:** Kirsty F. Houslay, Manuel van Gijsel-Bonnello, Tsvetana Petrova, Shaista Naqvi, J. Simon C. Arthur

## Abstract

IL-23 is an IL-12 family cytokine that is important in promoting Th17 responses and has been strongly linked to autoimmunity and psoriasis. It is a heterodimeric cytokine made up of a p19 subunit unique to IL-23 and a p40 subunit that is shared with IL-12. We show here that in response to LPS, the induction of IL-23p19 mRNA is regulated by a MSK1/2 – CREB dependent pathway downstream of ERK1/2 and p38α MAPK. Knockout of MSK1/2 resulted in a decrease in both IL-23p19 mRNA transcription and IL-23 secretion in GM-CSF differentiated bone marrow cells. Similar effects were seen when the MSK1/2 phosphorylation site in CREB was mutated to alanine. Stimulation with PGE2 promotes the nuclear localisation of CRTC3, a co-activator for CREB. In combination with LPS, PGE2 promoted IL-23p19 mRNA transcription and this was blocked by knockdown of CRTC3. Imiquimod induced skin inflammation in mice has been used as a model for psoriasis and is dependent on IL-23. While MSK1/2 knockout reduced the induction of IL-23 in vivo following i.p. injection of LPS, the knockout mice were not protected from Imiquimod induced skin inflammation. MSK1/2 knockout did not reduce the induction of IL-17 producing γδT cells following Imiquimod treatment, although MSK1/2 knockout did reduce the levels of these cells in mice receiving a control cream. The lack of protection in the Imiquimod model may be due to the known anti-inflammatory roles or MSKs, such as its contribution to the induction of IL-10.

## Introduction

Innate immune cells such as macrophages and dendritic cells respond to pathogens via Pattern Recognition Receptors (PRRs) that recognise specific Pathogen Associated Molecular Patterns (PAMPs). Toll like receptors (TLRs) comprise one such family of PRRs that play an important role in the host response to viral, microbial and fungal pathogens in both mice and humans. TLRs activate a number of intracellular signalling pathways, including Mitogen activated protein kinase (MAPK) cascades and the NFκB and IFR transcription factors[1-4]. Together these pathways induce the production of a range of pro-inflammatory molecules like cytokines and chemokines that help co-ordinate the inflammatory response to the pathogen. While inflammation is a key component of the host response, excessive inflammation can cause tissue damage and in extreme cases can lead to organ injury and mortality. It is therefore critical that inflammation is precisely regulated and a number of feedback pathways and anti-inflammatory mechanisms are deployed in order to achieve the right balance [5-7]. Paradoxically the same signalling pathways that promote inflammation are also involved in activating anti-inflammatory processes. One example of this is the p38α MAPK cascade, which promotes the production of the pro-inflammatory cytokine TNF by macrophages [8-10] and p38 inhibitors have been shown to reduce inflammation in mouse models of endotoxic shock [11, 12]. p38α however also serves to limit inflammation. For example, p38α phosphorylates and inhibits the TAB/TAK complex, which activates both NFκB and MAPK pathways following TLR activation [13]. p38α also helps promote the induction of DUSP1, a phosphatase that limits p38 and JNK activation [14-17], and the anti-inflammatory cytokine IL-10 [14].

Both IL-10 and DUSP1 can be regulated at transcriptional level in macrophages [15-19], and p38α has been described to regulate these genes via the activation of Mitogen and Stress Activated Protein Kinase (MSK) 1 and 2. MSK1 and 2 are nuclear protein kinases that can be activated downstream of either the ERK1/2 or p38α MAPK cascades [1]. For stimuli, such as TLR agonists, that activate both ERK1/2 and p38α, combined inhibition of both pathways is required to completely block MSK activation [20, 21]. A number of substrates have been described for MSKs including the transcription factors CREB and ATF1 and the chromatin protein histone H3 [22, 23]. Through phosphorylation of these substrates MSKs can regulate the induction of its target genes downstream of MAPK signalling. Both IL-10 and DUSP1 are CREB regulated, and CREB has been shown to be required for the maximal induction of these genes downstream of TLR agonists in macrophages [14, 21, 24-26]. MSKs phosphorylate CREB on Ser133 resulting in activation of its transcriptional activity [22, 27]. In addition to phosphorylation on Ser133, CREB can also be regulated by the CREB regulated transcription coactivator (CRTC) proteins. CRTCs can be phosphorylated by members of the Salt Inducible Kinase (SIK) family and these results in 14-3-3 binding and retention of the CRTC proteins in the cytoplasm [28]. Inhibition of SIKs allows the dephosphorylation of CRTC and their translocation to the nucleus where they can bind to CREB [28-30]. In macrophages, prostaglandin E2 (PGE2) has been shown to inhibit SIK mediated CRTC phosphorylation and promote CRTC3 localisation to the nucleus where it can help activate IL-10 transcription [29]. As a result, stimulation of macrophages with PGE2 and the TLR4 agonist LPS results in a greater production of IL-10 than stimulation with LPS alone.

In contrast to IL-10 knockout mice, which develop colitis [31], MSK1/2 double knockout mice are viable and do not develop any of the spontaneous phenotypes associated with IL-10 loss [22]. MSK1/2 knockout mice do however show an increased sensitivity to LPS induced endotoxic shock and produce higher levels of TNF, IL-6, IL-12 and IFNβ in response to LPS than wild type mice [21, 25, 32]. One of the plausible explanations for the differences in the phenotypes of MSK1/2 and IL-10 knockout mice is the possibility that MSKs and CREB may also regulate some pro-inflammatory genes in macrophages in addition to their described anti-inflammatory targets. In this study, we show that MSKs and CREB regulate the induction of IL-23p19 subunit downstream of TLR agonists. IL-23 is a member of the IL-12 family and has well established roles in the maintenance of Th17 cells and the development of autoimmunity [33-35]. Thus, like its upstream activator p38α, MSK1/2 can also induce both pro- and anti-inflammatory cytokines in innate immune cells.

## Results

### IL-23p19 mRNA induction is regulated via MSK – CREB pathway in macrophages

We have previously shown that PGE2 and LPS promote IL-10 induction via a CREB dependent mechanism in macrophages [29]. To identify other genes that might be regulated in a similar manner, we re-examined microarray data comparing bone marrow derived macrophages (BMDMs) stimulated with either LPS, PGE2 or a combination of LPS and PGE2 for 1 h. 139 genes were identified as being upregulated more than 4-fold by LPS alone. 18 of these genes were upregulated further 2-fold when a combination of LPS and PGE2 was used (Fig 1A); of these *Il10, IL-6, Ptgs2, NR4A1, NR4A3, Trem1, Gja1* and *IL1rn* have previously been reported as CREB or MSK regulated genes [36-42]. In addition, *Il23a* was identified in this list. *Il23a* encodes the p19 subunit of IL-23, a pro-inflammatory cytokine implicated in the maintenance of IL-17 producing T cells [43, 44]. *Il23a* was selected for further study as *Il23a* knockout in mice has been shown to be protective in a range of autoimmune models including experimental autoimmune encephalomyelitis (EAE), arthritis and imiquimod-induced skin inflammation [44-46]. To confirm the microarray data, IL-23p19 mRNA was analysed by qPCR in independent samples. As expected, this showed a similar pattern of IL-23p19 mRNA regulation to the array data (Fig 1B).

**Figure 1.**
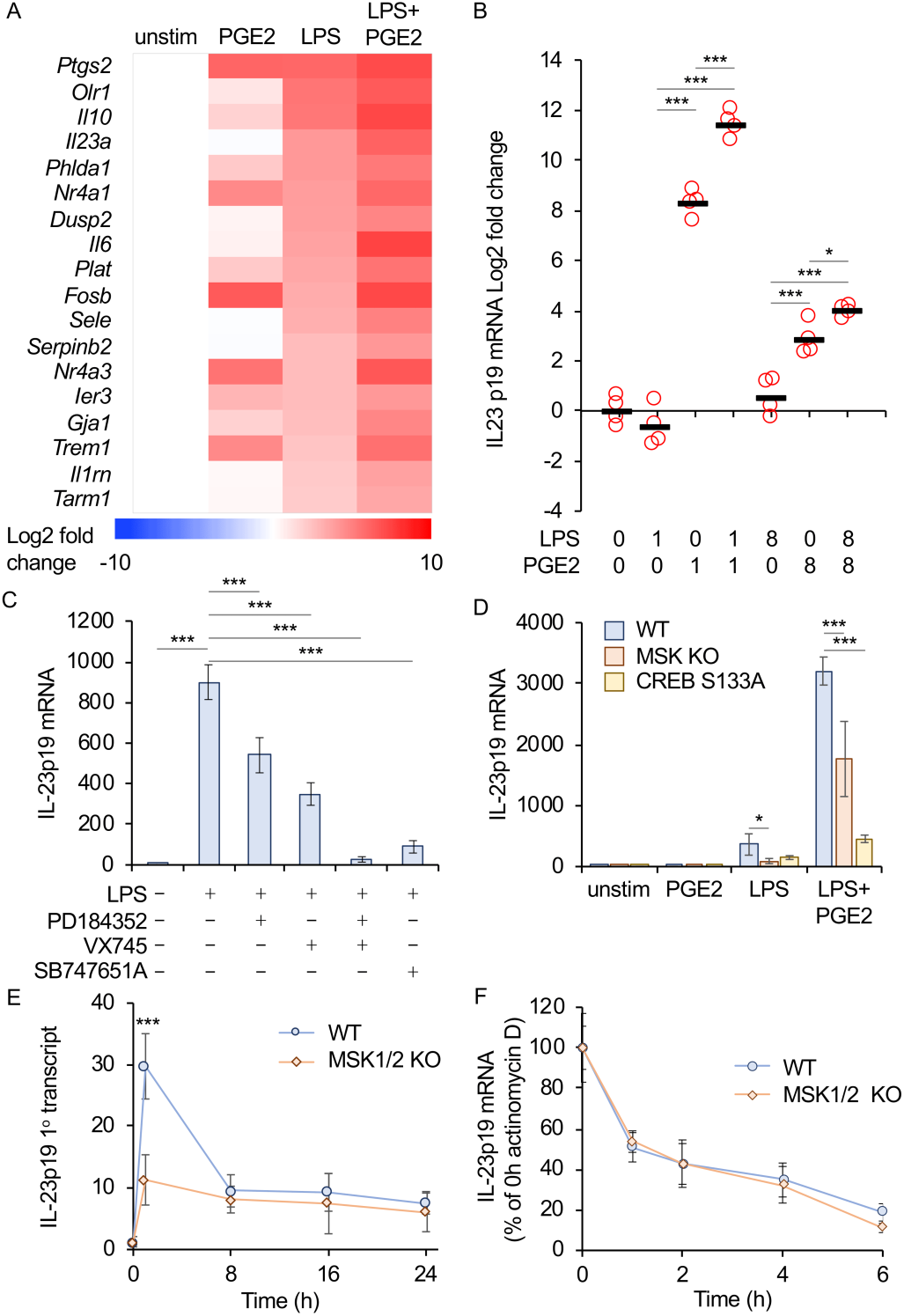
MSK1/2 and CREB regulate IL-12p19 mRNA induction in macrophages. A) Heat map of genes that were upregulated 4 fold in BMDMs by 100 ng/ml LPS for 1 h and a further 2 fold when a combination of LPS and 10 μM PGE2 was used. B) BMDMs were stimulated for 1 or 8h with 100 ng/ml LPS and/or 10 μM PGE2 as indicated. Total mRNA was isolated and IL-23p19 mRNA levels determined by qPCR. Bars show average value and circles represent individual values from BMDMs from separate mice. Within time points, a *p*<0.05 is indicated by * and *p*<0.001 by *** (2 way ANOVA and Holm-Sidak post hoc testing). C) Where indicated BMDMs were treated with 2 μM PD184352, 1 μM VX745 or 10 μM SB747651A before stimulation with 100 ng/ml LPS for 1 h. IL-23p19 mRNA levels were determined by qPCR. Compared to the LPS condition, a *p*<0.001 is indicated by *** (1 way ANOVA and Holm-Sidak post hoc testing) D) BMDMs were isolated form wild type, MSK1/2 knockout and CREB S133A knockin mice. Cells were stimulated for 1 h with either 10 μM PGE2, 100 ng/ml LPS or a combination of LPS and PGE2. IL-23p19 mRNA levels were measured by qPCR. For comparisons to the wild type stimulations, a *p*<0.05 is indicated by * and *p*<0.001 by *** (2 way ANOVA and Holm-Sidak post hoc testing). E) Wild type or MSK1/2 knockout BMDMs were stimulated for the indicated times with 100ng/ml LPS and IL-23p19 primary transcript levels determined by qPCR. Between genotypes *** indicates *p*<0.001 (2 way ANOVA and Holm-Sidak post hoc testing). F) Wild type or MSK1/2 knockout BMDMs were stimulated for 1h with 100ng/ml LPS before addition of 1 μg/ml actinomycin D. IL-23p19 mRNA levels were determined at 0, 1, 2, 4 and 6 h after actinomycin D addition. Wild type and MSK1/2 knockouts were not significantly different (2 way ANOVA). In B-E mRNA levels are expressed as fold induction relative to unstimulated wild type cells. In C to F, graphs show the average and standard deviations of independent cultures from 4 mice per genotype.

To determine if IL-23p19 mRNA induction was regulated via an MSK – CREB dependent pathway, a combination of small molecule inhibitors and mouse genetics was used. Following LPS stimulation of macrophages, the ERK1/2 and p38 MAPK pathways activate MSK1/2 which phosphorylate CREB at S133 [21]. Inhibition of the ERK1/2 pathway with PD184352 or the p38 pathways with VX745 reduced IL-23p19 mRNA induction, while a combination of both inhibitors further reduced IL-23p19 mRNA induction in response to LPS. The induction of IL-23p19 mRNA was also decreased by the MSK inhibitor SB-747651A (Fig 1C). To confirm these results, Bone Marrow-derived Macrophages (BMDMs) were prepared from wild type, MSK1/2 double knockout and CREB-S133A knockin mice (Fig 1D). The induction of IL-23p19 mRNA was decreased in both the MSK1/2 knockout and CREB S133A knockin BMDMs relative to wild type cells in response to LPS. PGE2 did not induce IL-23p19 mRNA in any of the genotypes, demonstrating a role for an MSK – CREB dependent pathway in LPS induced IL-23p19 mRNA induction. MSK1/2 knockout or CREB S133A knockin BMDMs showed reduced IL-23p19 mRNA induction relative to wild type cells in response to a combination of LPS and PGE2. A combination of LPS and PGE2 gave a stronger induction of IL-23p19 mRNA than LPS alone for all 3 genotypes, suggesting that the effect of PGE2 is independent of MSK1/2 and CREB S133 phosphorylation (Fig 1D). Changes in mRNA levels can be due to changes in either transcription or mRNA stability. Changes in the level of the primary RNA transcript for a gene can provide a better readout for changes in transcription than analysis of mRNA levels. We therefore looked at the effect of MSK1/2 knockout on IL-23p19 transcript levels following LPS stimulation. As for the mRNA, MSK1/2 knockout reduced IL-23p19 primary transcript levels (Fig 1E). To determine if MSKs affected IL-23p19 mRNA stability, BMDMs were treated with LPS for 1h and then actinomycin D was added to block further transcription. The loss of IL-23p19 mRNA was then measured over time showing no effect on IL-23p19 mRNA stability in MSK1/2 knockout compared to wildtype BMDMs (Fig 1F). Together these results would indicate that MSK1/2 regulate IL-23p19 mRNA transcription via CREB phosphorylation. PGE2 activates the cAMP-PKA pathway in macrophages, which is able to promote the nuclear localisation of the CRTC family of CREB co-activator proteins [29]. We have previously shown that CRTC3 is important for the regulation of IL-10 in macrophages using siRNA [29]. We therefore analysed the effect on IL-23p19 mRNA induction in these experiments. Knockdown of CRTC1 or 2 did not affect IL-23p19 mRNA induction. In contrast, while CRTC3 knockdown did not affect LPS-induced IL-23p19 mRNA induction, it did block the increase seen when PGE2 was added in combination with LPS (Fig 2A). Successful knockdown of the respective CRTC isoforms was confirmed by qPCR (Fig 2B). CRTC3 is phosphorylated by a member of the SIK family causing the CRTC3 to be retained in the cytoplasm. Activation of PKA downstream of PGE2 allows PKA to phosphorylate SIKs and block their ability to phosphorylate CRTC3 [29]. In agreement with this, treatment with MRT67307, which inhibits all three SIK isoforms, mimicked the effect of PGE2 on LPS stimulated IL-23p19 mRNA induction (Fig 2C).

**Figure 2.**
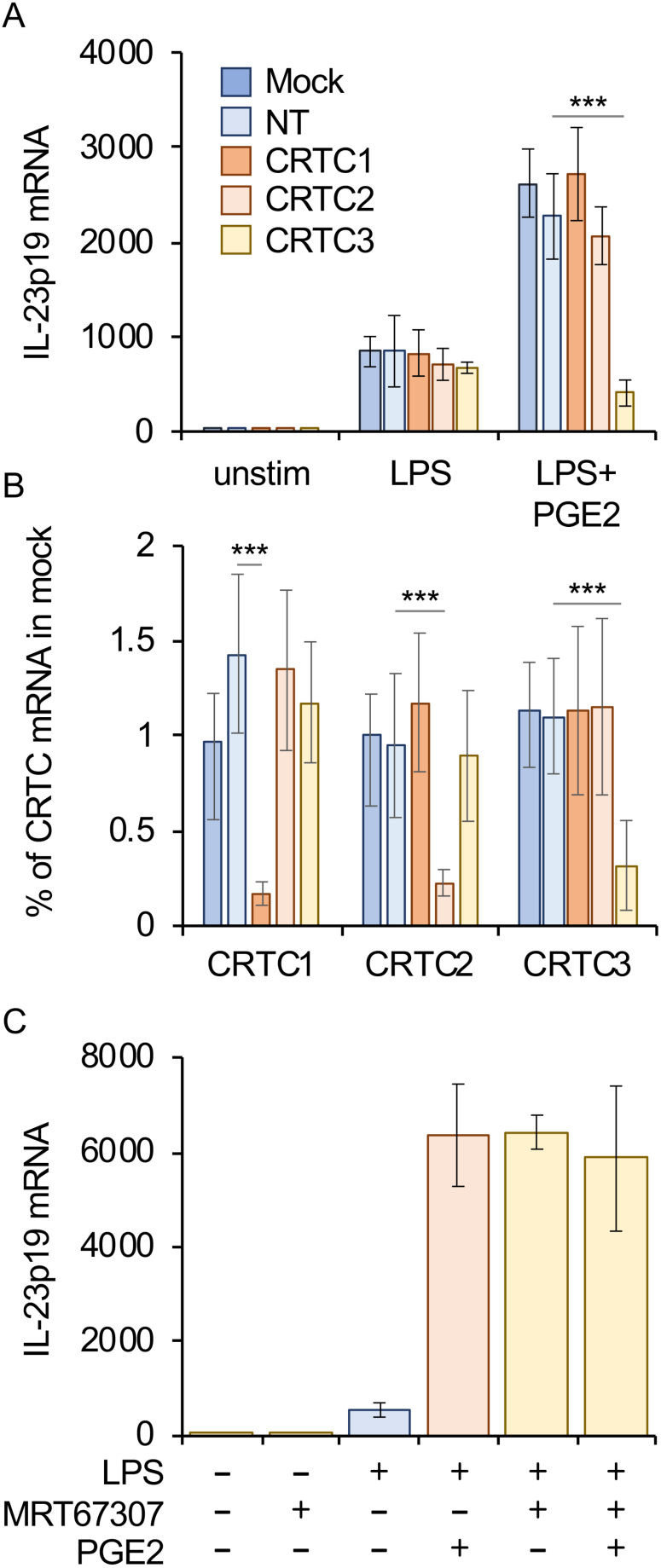
PGE2 promotes IL-23p19 mRNA induction via CRTC3. A) Wild-type BMDMs were transfected with siRNA against CRTC1, 2, or 3, a nontargeting siRNA control (NT) or no siRNA (mock). Twenty-four to 48 hours after transfection cells were stimulated with either 100 ng/ ml LPS or LPS and 10 μM PGE2 for 1 h, and the levels of IL-23p19 mRNA were determined by qPCR. B) As (A) but mRNA levels for CRTC1, CRTC2 and CRTC3 were determined. Values for the unstimulated conditions are shown but similar results were obtained for the 1h time point. C) Where indicated wild type BMDMs were incubated for 1 h with 2 μM MRT67307 before stimulation with 100ng/ml LPS and/or 10 μM PGE2 for a further hour. IL-23p19 mRNA levels were then determined by qPCR. Graphs show the average and standard deviation of culture from 4 mice. In A and B, for comparisons to the non-targeting siRNA, a p<0.001 is indicated by *** (2 way ANOVA and Holm-Sidak post hoc testing).

### MSKs regulate the production and secretion of IL-23

IL-23 is a heterodimer of IL-23p19 and IL-12p40 subunits. While BMDMs upregulate both p19 and p40 mRNA in response to LPS they do not secrete measurable amount of IL-23 [47]. In contrast, bone marrow cells differentiated in GM-CSF (GM-BMCs), which gives rise to heterogenous population of macrophages and dendritic cells [48], do secrete detectable levels of IL-23 in response to TLR stimulation [47]. We first confirmed that MSKs were activated in GM-BMCs in response to LPS (Fig 3A). TLR stimulation activated ERK1/2 and p38, as judged by phosphorylation on the TXY motifs. In line with this LPS stimulated the phosphorylation of MSK1 on Thr581, a site that correlates with MSK1 activation [20]. LPS stimulation also resulted in the phosphorylation of the MSK substrate CREB, and this phosphorylation was absent in GM-BMCs from MSK1/2 knockout mice (Fig 3A). LPS increased the levels of IL-23p19 mRNA, as well as the mRNA for IL-12p40 and IL-12p35 (Fig 3B). As in MSK1/ 2 knockout GM-BMCs, the induction of IL-23p19 mRNA was reduced relative to wild type GM-BMCs (Fig 3B). The levels of IL-12p40 were not affected by MSK1/2 knockout at most of the time points analysed, while IL-12p35 mRNA was increased at later time points (Fig 3C and D). This increase in IL-12p35 may be due to the ability of MSKs to regulate IL-10 induction in GM-BMCs [25], as IL-10 can set up a negative feedback loop to repress IL-12p35 induction. Consistent with this, MSK1/2 knockout reduced IL-10 secretion in response to LPS and increased IL-12p70 secretion (Supplementary Fig 1A). Similar to the findings in BMDMs, PGE2 in combination with LPS resulted in a higher induction of IL-23p19 mRNA than LPS alone (Fig 3E and F). Relative to wild type GM-BMCs, both MSK1/2 knockout and CREB S133A knockin reduced the induction of IL-23p19 mRNA in response to either LPS or LPS and PGE2 in combination. LPS and PGE2 in combination gave a higher induction of IL-23p19 mRNA than LPS alone in all 3 genotypes, suggesting that the effect of PGE2 did not require CREB S133 phosphorylation (Fig 3E and F).

**Figure 3.**
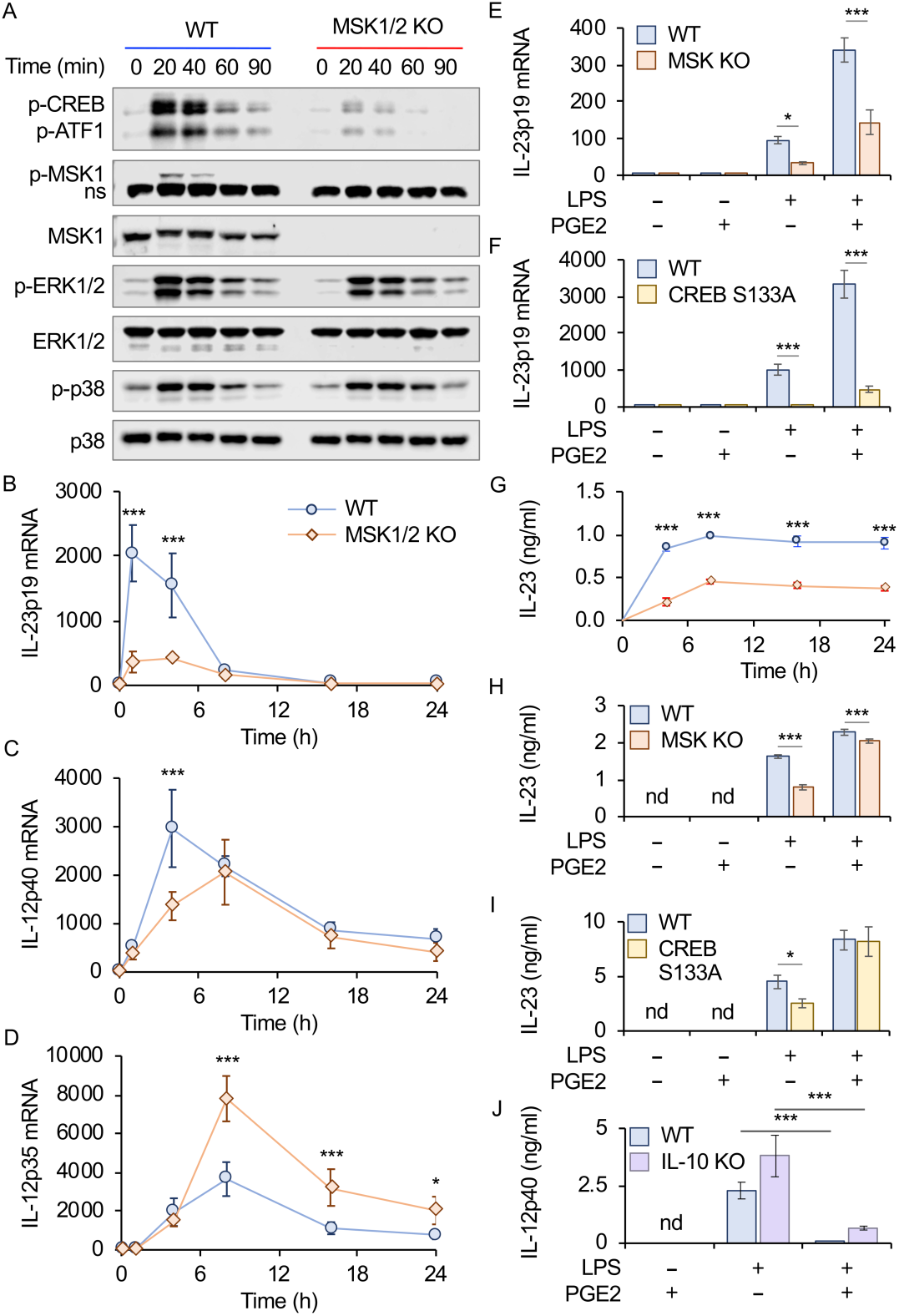
Regulation of IL-23 production by MSK1/2 and CREB in GM-BMCs. A) Wild type or MSK1/2 knockout GM-BMCs were stimulated for the indicated times with 100 ng/ml LPS and the levels of the indicated phospho and total proteins determined by immunoblotting. ns indicates a non-specific band. B-D) Wild type or MSK1/2 knockout GM-BMCs were stimulated for the indicated times with 100 ng/ml LPS and the levels of IL-23p19 (B), IL-12p40 (C) and IL-12p35 (D) determined by qPCR. Wild type or MSK1/2 knockout GM-BMCs were stimulated for 1 h with 100 ng/ml LPS and/or 10 μM PGE2 as indicated and IL-23p19 mRNA levels determined. F) Wild type or CREB S133A knockin GM-BMCs were stimulated for 1 h with 100 ng/ml LPS and/or 10 μM PGE2 as indicated and IL-23p19 mRNA levels determined. G) Wild type or MSK1/2 knockout GM-BMCs were stimulated for the indicated times with 100 ng/ml LPS and the level of IL-23 secreted determined by ELISA. H) Wild type or MSK1/2 knockout GM-BMCs were stimulated for 8 h with 100 ng/ml LPS and/or 10 μM PGE2 as indicated and the level of IL-23 secreted determined by ELISA. I) Wild type or MSK1/2 knockout GM-BMCs were stimulated for 8 h with 100 ng/ml LPS and/or 10 μM PGE2 as indicated and the level of IL-23 secreted determined by Luminex assay. J) Wild type or IL-10 knockout GM-BMCs were stimulated with 100 ng/ml LPS or LPS and 10 μM PGE2 for 8h and IL-12p40 secretion measured by a Luminex based assay. In B to J data represent average and standard deviation from BMDMs from 4 mice per genotype. nd indicates conditions were the level of cytokine was below what could be detected in the assay. In B to I, for comparisons between wild type and MSK1/2 a p<0.001 in shown by *** (B-D 1 way ANOVA, E-I 2 way ANOVA with Holm-Sidak post hoc testing). In J a difference between the LPS and LPS + PGE2 condition with a p<0.001 (Students ttest) in indicated by ***.

In line with the reduction in IL-23p19 mRNA, MSK1/2 knockout reduced the secretion of IL-23 following LPS stimulation (Fig 3G and H). As has previously been reported, treatment of BMDMs with GM-CSF resulted in the detectable secretion of IL-23 following LPS stimulation (Supplementary Fig 1B, [47]). Under these conditions MSK1/2 also resulted in a reduced secretion of IL-23 relative to with type GM-BMCs (Supplementary Fig 1B). In agreement with the data for MSK1/2 knockouts, CREB S133A knockin also reduced IL-23 secretion in response to LPS in GM-BMCs (Fig 3I). Treatment with LPS and PGE2 together resulted in a modest increase in IL-12 production relative to IL-23 alone (Fig 3H and I). Surprisingly, in response to LPS and PGE2 neither MSK1/2 knockout or CREB S133A knockin reduced IL-23 secretion relative to wild type cells (Fig H and I), despite IL-23 mRNA being decreased (Fig E and F). One potential reason is that IL-23 is a heterodimer of IL-23p19 and IL-12p40. In response to LPS alone, IL-12p40 production may not be limiting. However, while PGE2 promotes IL-23p19 production, it strongly inhibits the production of IL-12p40 via both IL-10 dependent and independent mechanisms (Fig 3J).

As MSK1/2 knockout decreased IL-23 secretion from isolated GM-BMCs, we next determined if MSKs were required for optimal IL-23 production *in vivo*. Mice were given an i.p. injection of either LPS or PBS and IL-23 levels measured in the serum. Wild type mice produced IL-23 in response to LPS. In line with the experiments in GM-BMCs, MSK1/2 knockout mice produced significantly less IL-23 following LPS injection than wild type animals (Fig 4)

**Figure 4.**
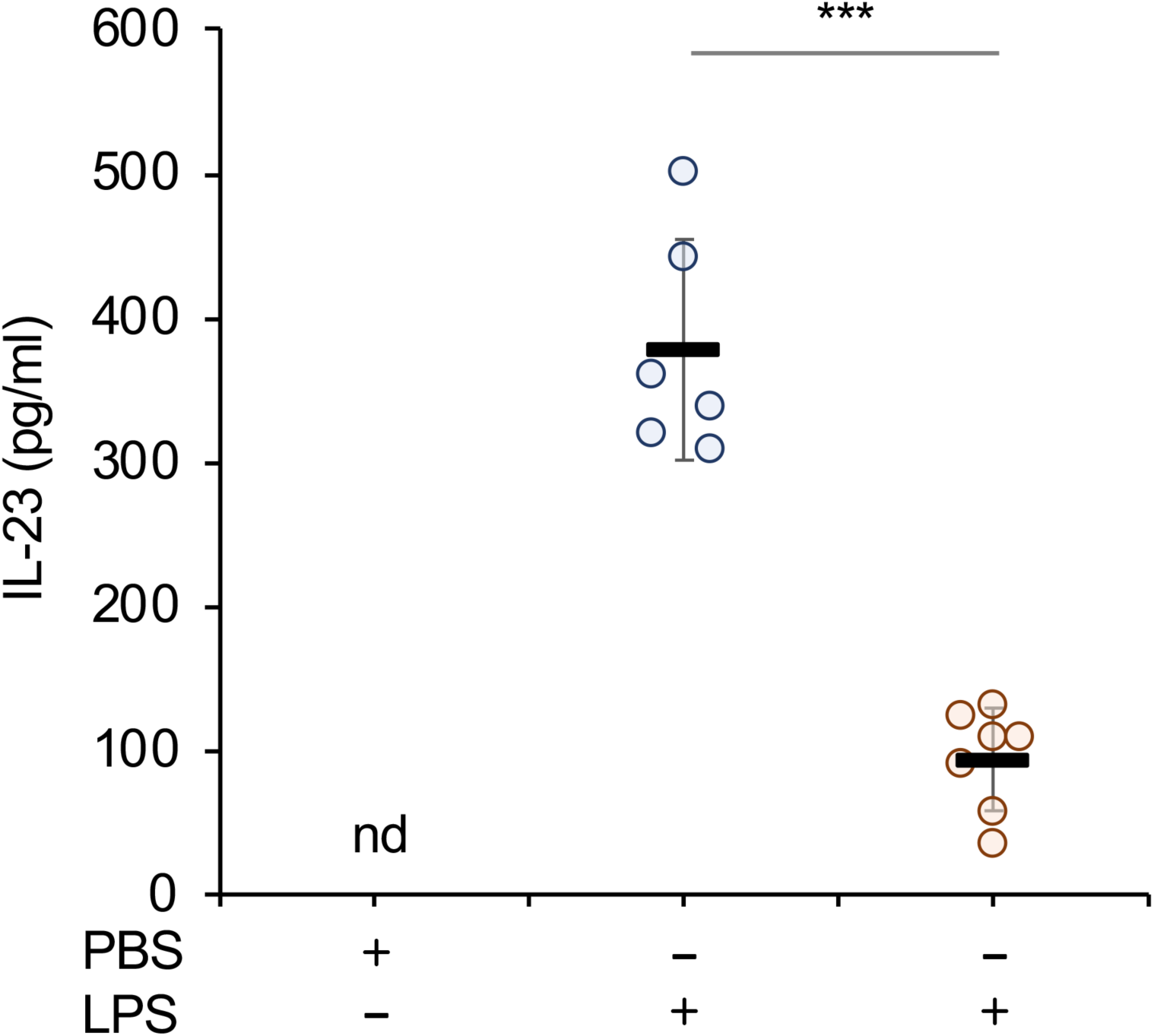
MSK1/2 regulate IL-23 induction following i.p. injection of LPS. Wild type or MSK1/2 knockout mice were given an i.p. injection of either LPS (2mg/kg) or PBS vehicle as indicated. After 4 h serum levels of IL-23 were measured by ELISA. Bars show average and standard deviation with measurements from individual animals shown by circles. Relative to wild type LPS injected mice a *p* value (two tailed Students ttest) of < 0.001 is shown by ***.

### Effect of MSK1/2 knockout on *in vivo* IL-23 production and imiquimod induced skin inflammation

In mice, treatment of the skin with Aldara, a cream containing the TLR7 agonist imiquimod, induces a psoriasis like skin inflammation. Knockout of the IL-23p19 subunit has been shown to protect mice from inflammation in this model [45]. The ears of wild type and MSK1/2 knockout mice were treated with Aldara or Oilatum (a control cream that does not contain imiquimod) for 7 days after which mice were untreated for 4 days to allow inflammation to resolve. Ear thickness was measured throughout this period as measurement of inflammation. Thickening was observed in both wild type and MSK1/2 knockout mice treated with Aldara, and in both cases this started to resolve once treatment was stopped. No differences were however observed between the two genotypes in the changes in ear thickness (Fig 5A). Analysis of IL-17 levels in biopsies taken on day 5 and day 11 showed that IL-17A and IL-17F levels were induced in the ear by Aldara treatment, however again there were no significant differences between the wild type and MSK1/2 knockout mice (Fig 5B and C). In agreement with this, IL17A and F mRNA levels in ear biopsies were not affected by MSK1/2 knockout (Fig 5D and E). Increased levels of IL-23p19 mRNA were detected following Aldara treatment in wild type mice, however in contrast to the previous results in GM-BMCs, the levels were not affected by MSK1/2 knockout (Fig 5F) while IL-12p40 mRNA levels were also not affected (Fig 5G).

**Figure 5.**
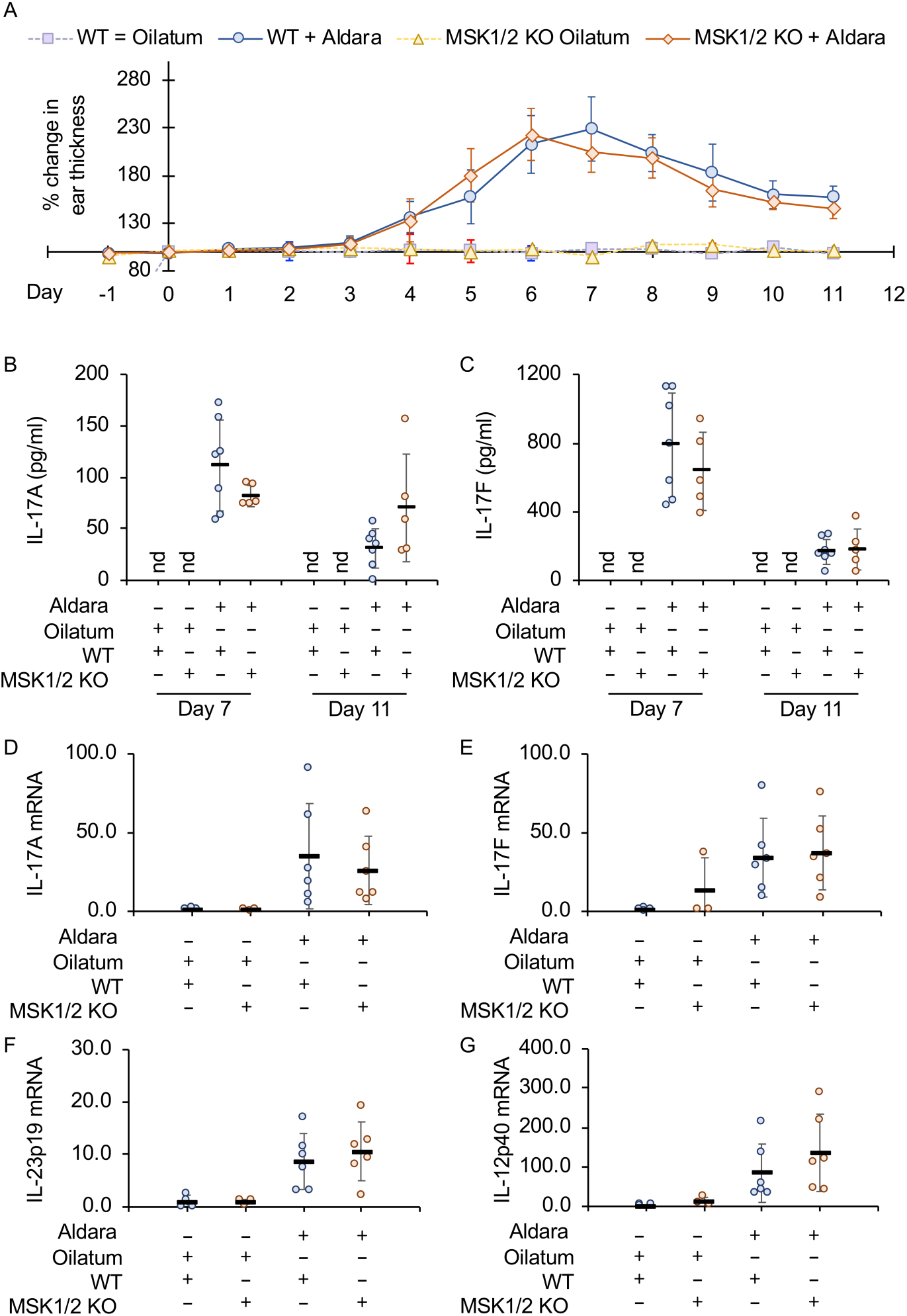
Imiquimod induced skin inflammation in MSK1/2 knockout mice. A-C) Wild type or MSK1/2 knockout mice were treated daily from day 1 to 7 with either Aldara or Oilatum cream as described in the methods (WT Oilatum n=5, MSK1/2 KO Oliatum n=2, Wild type Aldara n=7, MSK1/2 KO Aldara n=5). Ear thickness is shown in (A) and IL-17A levels in ear biopsies in (B) and IL-17F levels in (C). D-G) Wild type or MSK1/2 knockout mice were treated daily with Aldara or Oilatum cream for 6 days (WT Oilatum n=4, MSK1/2 KO Oliatum n=3, Wild type Aldara n=6, MSK1/2 KO Aldara n=6). RNA was then isolated from the ear and levels of IL-17A (D), IL-17F (E), IL-23p19 (F) and IL-12p40 (G) mRNA determined. Values show fold change relative to the average values of wild type Oilatum treated mice. Graphs show mean and standard deviation with individual mice in B to F shown by circles. No effect of genotype was observed (p>0.05, 3 way ANOVA for A, two tailed students ttest for B-G),

In addition to promoting inflammation in the ear, Aldara also has systematic effects in the mice. Mice treated with Aldara for 6 days showed an increase in spleen size as well as increased numbers of neutrophils, Ly6C+ve and Ly6C-ve monocytes in the spleen (Fig 6A to E, Supplementary Fig 2). The numbers of neutrophils and monocytes in the spleens of Aldara treated mice was not affected by knockout of MSK1 and 2. There was however a slight decrease in the numbers of neutrophils in MSK1/2 knockout mice compared to wild type animals receiving the control Oilatum cream. Previous studies have shown that IL-23 is involved in the induction of IL-17 producing γδ-T cells following Aldara treatment [49, 50]. The numbers of γδ T cells were similar between wild type and MSK1/2 knockout mice (Supplementary Fig 3). Aldara increased the proportion of γδ-T cells in spleen able to make IL-17 on ex vivo stimulation with PMA, however this was not affected by MSK1/2 knockout (Fig 6F). In mice treated with Oilatum the percentage of T cells able to produce IL-17 was lower in MSK1/2 knockout mice relative to wild types (Fig 6F), and a similar result was found in the lymph nodes (Supplementary Fig 3).

**Figure 6.**
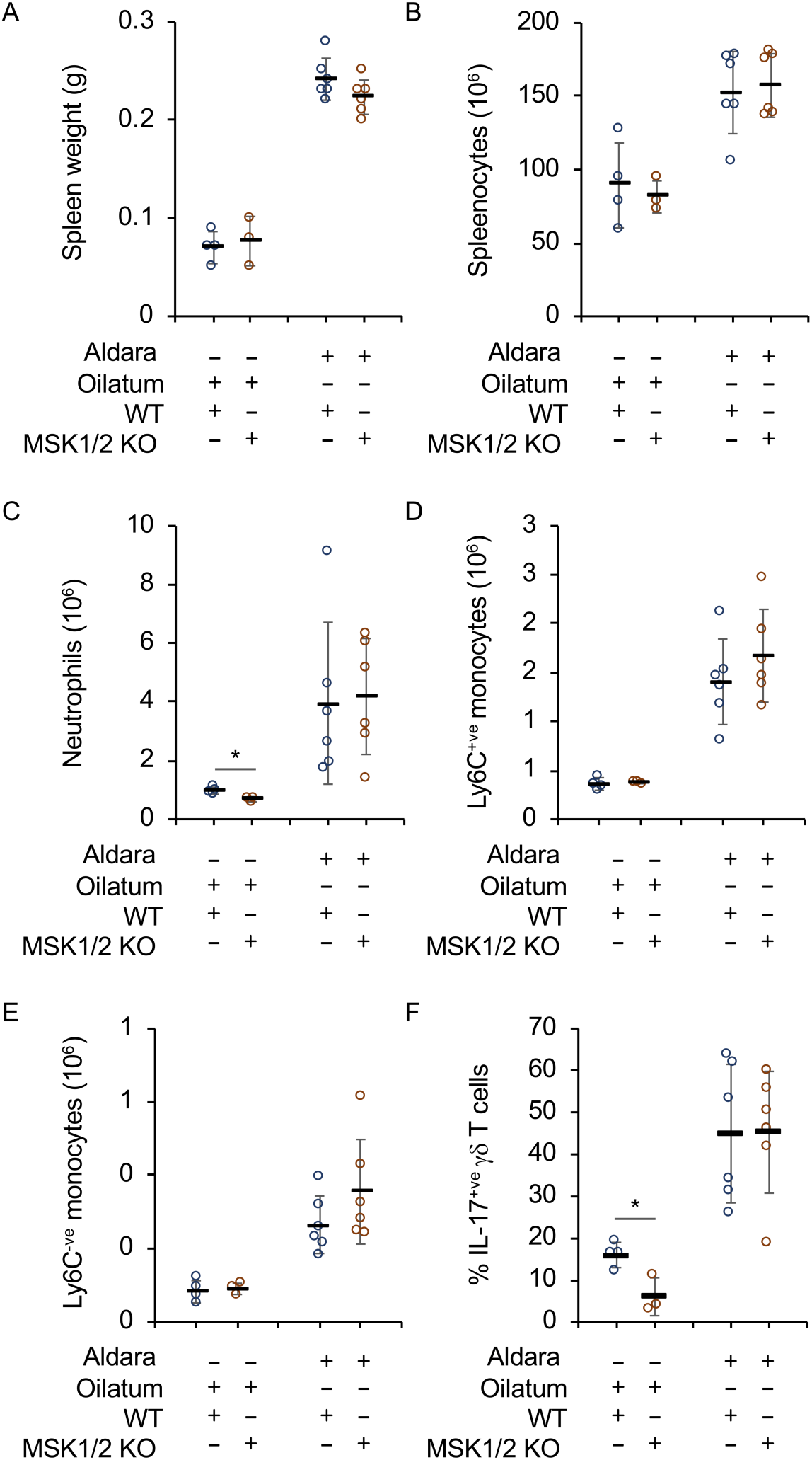
Effect of imiquimod on myeloid cells and γδ T cells. Wild type or MSK1/2 knockout mice were treated daily with Aldara or Oilatum cream for 6 days (WT Oilatum n=4, MSK1/2 KO Oliatum n=3, Wild type Aldara n=6, MSK1/2 KO Aldara n=6). Spleens were isolated and analysed by flow cytometry as described in the methods. Graphs show spleen weight (A), total number of splenocytes following lysis of red blood cells (B), numbers of neutrophils (C), Ly6C^+ve^ monocytes (D), Ly6C^-ve^ monocytes (E). In (F), cells were stimulated with ionomycin and PMA and the percentage of γδ T cells expressing IL-17 quantified by flow cytometry as described in the methods. Graphs show mean and standard deviation with individual mice in B to F shown by circles. Between wild type and MSK1/2 knockouts, a p<0.05 in indicated by * (two tailed Students ttest).

## Discussion

In this study, we show that in BMDMs and GM-BMCs, the induction of IL-23p19 mRNA downstream of TLR4 requires the activation of an MSK1/2 – CREB dependent pathway (Fig 7). MSKs are activated via the ERK1/2 and p38 MAPK pathways [20, 22]. Therefore, the role for MSK1/ 2 in regulating IL-23 production is consistent with the findings that ERK1/2 and/or p38α pathways are involved in this process [50-52]. For example, inhibition of ERK1/2 or p38 was reported to reduce the induction of a IL-23p19 luciferase reporter in macrophages [53] while inhibition of ERK1/2 and or p38 reduced IL-23p19 induction in Langerhans cells, epithelial cells and colorectal cancer cells.

**Figure 7.**
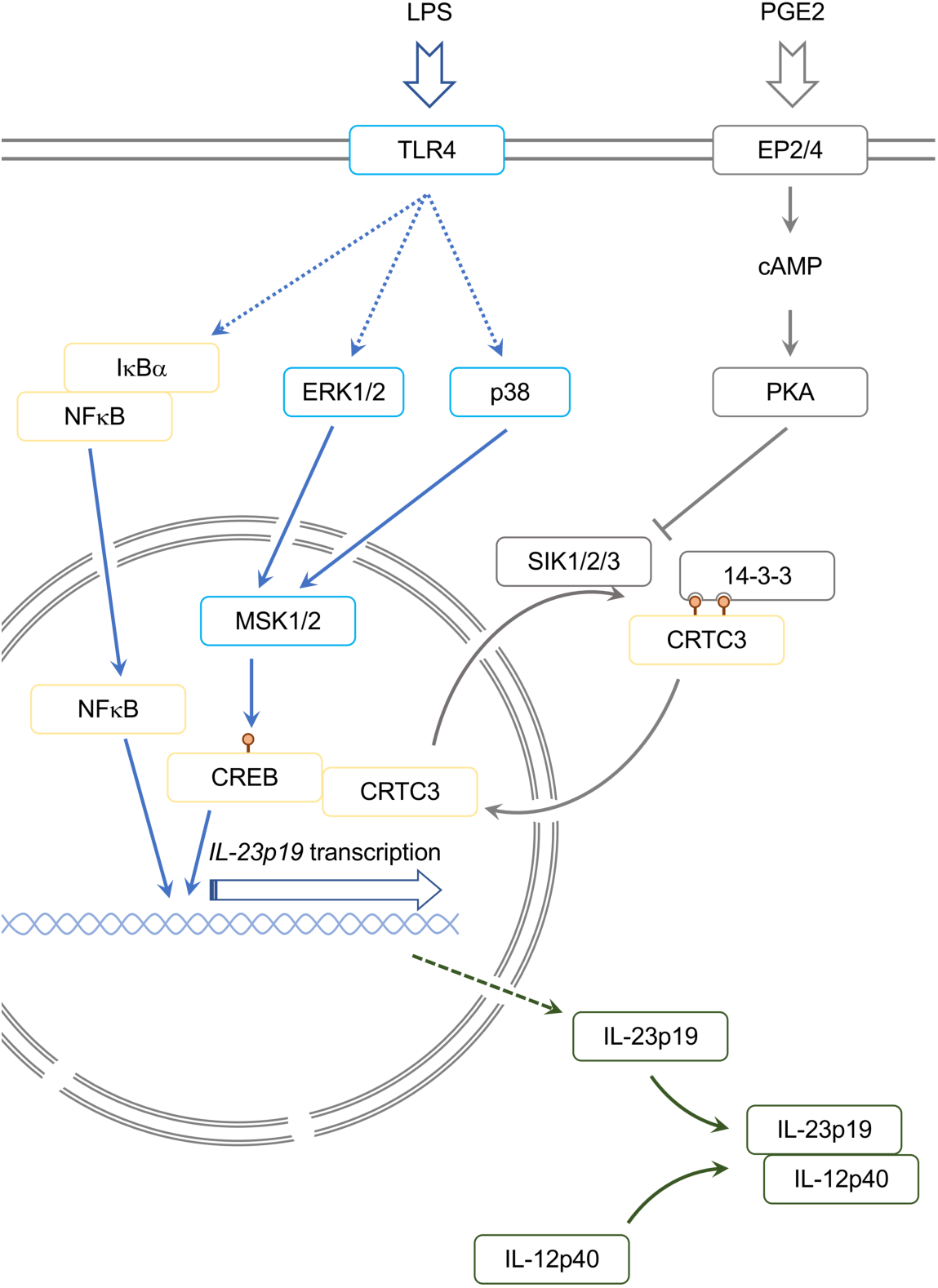
Model for control of IL-23p19 induction. LPS acts via TLR4 to stimulate the activation of the ERK1/2 and p38 MAPK pathways, leading to the activation of MSK1 and 2. MSKs then phosphorylate CREB on Ser133 on the IL-23p19 promoter, leading to the activation of IL-23p19 transcription. CREB phosphorylation alone is insufficient to activate the IL-23p19 promoter, in addition TLR4 activation results in the degradation of IκBα leading to the release and nuclear translocation of NFκB. Once in the nucleus NFκB bind the IL-23p19 promoter and in combination with CREB S133 phosphorylation this results in IL-23p19 transcription. CRTC3 can act as a co-activator for CREB. In the absence of stimulation CRTC3 is phosphorylated by one or more of the SIK isoforms. This creates a binding site for 14-3-3 proteins and the retention of CRTC3 in the cytoplasm. PGE2 acts via the EP2 and ER4 receptors to stimulate the production of cAMP which activates PKA. PKA then phosphorylates SIKs which inhibits the ability of SIKS to phosphorylate CRTC3 in the cells. This allows CRTC3 dephosphorylation, resulting in its translocation to the nucleus where it acts as a CREB co-activator at the IL-23p19 promoter. Once transcribed, IL-23p19 protein heterodimerises with IL-12p40 to form IL-23 which can then be secreted from the cell.

MSK1/2 knockout has been found to sensitise mice to LPS induced endotoxic shock [21]. This is partly due to a reduction in the production of the anti-inflammatory cytokine IL-10, accompanied by an increased production of a number of pro-inflammatory cytokines including TNF, IL-6 IL-12 and IFNβ [21, 26, 32]. The ability of MSKs to promote IL-23p19 induction is however unlikely to be dependent on its role in controlling IL-10 production, as IL-10 represses IL-23p19 mRNA induction in response to LPS ([53] Supplementary Fig 1C). Although the majority of previously described MSK targets in macrophages, such as IL-10, IL-1ra, DUSP1 and TTP, all have anti-inflammatory roles, MSK1/2 knockout mice do not show overt signs of spontaneous inflammatory pathologies. One reason for this may be that the anti-inflammatory role of MSKs is cell type restricted to a subset of immune cells; for example while MSK1/2 regulates LPS induced IL-10 production in macrophages it is not required for IL-10 secretion by LPS stimulated B cells [21, 54]. An additional possibility is that MSKs have some pro-inflammatory roles in addition to their known anti-inflammatory functions. The regulation of IL-23 production by MSK1/2 would be one such pro-inflammatory function. In this respect, MSKs over all effects may have similarities to those of p38α. While p38α activation was originally thought of as pro-inflammatory signal, further work has shown that p38α is able to activate both pro and anti-inflammatory pathways in innate immune cells [13, 14]. p38α activates the kinases MK2 and 3 in addition to MSK1 and 2. Given that MK2/3 are important in the production of TNF, while MSK1/2 regulate IL-10 it has been suggested that substrate selective p38α inhibitors or MK2/3 inhibitors may retain the ability to block pro-inflammatory cytokines while sparing IL-10 production [55]. This has not proved true – we show here that MSKs are required for IL-23 production and other studies have found that MK2/3 are required for IL-10 production in addition to regulating TNF [55, 56]. Thus both MSKs and MKs play both pro and anti-inflammatory roles downstream of p38α.

While LPS stimulates CREB phosphorylation via MSK1/ 2, PGE2 activates PKA which can also phosphorylate CREB on S133 in macrophages [29]. Despite this, PGE2 alone did not induce a strong induction of IL-23p19 mRNA. This may be due to a requirement for additional LPS activated transcription factor binding sites on the IL-23p19 promoter that may be not be activated by PGE2 alone. For example, several putative NFκB sites have been identified and mutation of the two proximal sites has been shown to reduce transcription from the IL-23p19 promoter. Consistent with this, knockout of the c-Rel NFκB subunit reduces LPS stimulated IL-23p19 mRNA induction in both macrophages and dendritic cells [57, 58]. An ability of PGE2 to enhance LPS and β-glucan stimulated IL-23 production in dendritic cells has been noted previously [59-61]. While CREB is required for the effects of PGE2 on IL-23 [59, 62], CREB phosphorylation is not critical and instead the effect is mediated via a SIK – CRTC3 dependent pathway (Fig 1-2). The ability of PGE2 to increase TLR induced IL-23p19 mRNA transcription is in contrast to its inhibitory effects on the induction of IL-12p40, which is a second component in the IL-23 heterodimer. While the effects of PGE2 on IL-23p19 transcription are clear, its effects on IL-23 secretion are more complex as IL-12p40 may become limiting in the formation of the IL-23 heterodimer. While in dendritic cells PGE2 generally enhances IL-23 secretion, in human monocytes PGE2 inhibits IL-23 secretion as the IL-12p40 subunit is limiting [63]. This may explain why MSK1/2 knockout affected IL-23p19 mRNA induction following LPS and PGE2 stimulation when it did not have a significant effect of IL-23 secretion (Fig 3).

IL-23 is important for the pathogenesis of psoriasis and IL-23 neutralising antibodies have proved successful in the clinic for psoriasis treatment [64-66]. Increased phosphorylation of both MSK1 and CREB have been reported in psoriatic skin lesions, suggesting potential role for MSKs in the disease[67-69]. Imiquimod induced skin inflammation has been used as a mouse model for psoriasis [65] and knockout of IL-23p19 is protective in this model [45].

While MSK1/2 reduced IL-23 production from cultured GM-BMCs and following i.p. injection of LPS in mice, it did not have a major impact on imiquimod induced skin inflammation. This may be due to the complex effects of MSKs on innate immunity. Although MSK1/2 knockout reduces IL-23 production in response to LPS, it also reduces IL-10 production while promoting the production of TNF, IL-6, IL-12, INFβ and PGE2 by TLR stimulated macrophages. Related to this, knockout of IL-10 mice results in increased inflammation in imiquimod induced skin inflammation as well as elevated levels of IL-23p19 mRNA [70]. Interestingly, while MSK1/2 knockout did not affect the numbers of IL-17 producing γδ T cells in the spleens of Aldara treated mice, they were reduced in Oilatum treated animals. This could suggest that the Aldara treatment saturates the response and a difference is only seen when the potential inflammatory stimulus is much weaker.

In summary, we show that IL-23 induction in cultured cells can be regulated via an MSK – CREB dependent pathways downstream of TLR activation. Despite this MSK1/2 knockout did not prevent the development of imiquimod induced skin inflammation, a process known to be dependent of IL-23, suggesting a more complex effect of MSK1/2 knockout *in vivo*.

## Methods

### Mice

MSK1 and MSK2 knockout and CREB S133A knockin mice have been described previously [22, 71]. MSK1/2 mice were backcrossed onto C57Bl/6 (>18 generations). CREB mice had been backcrossed to C57Bl/6 for a least 12 generations. Non breeding mice were housed in same-sex groups, in individually ventilated sterile cages and were given standard diet R&M1 (SDS, Special Diets Services). Animals were maintained in rooms with controlled 12□h/12□h light/dark cycle, 21□°C temperature, and relative humidity of 45–65%. All the work was performed under a UK Home Office project license in accordance with UK and EU regulations and subject to local ethical review by the University of Dundee Ethical Review Committee.

To measure *in vivo* cytokine induction, mice were given an i.p. injection of either PBS or 2 mg/kg LPS (Sigma (E. coli 055:B5)) dissolved in PBS. After the indicated time, mice were culled and blood taken by cardiac puncture.

### Array datasets

A microarray data set on unstimulated BMDMs and BMDMs treated with LPS, PGE2 or LPS + PGE_2_ for 1 h using Affymetrix mouse gene 1.1 ST arrays has been described previously [29] and has the accession number GSE41833.

### Cell culture

BMDMs and GM-BMCs were cultured as described [72]. Briefly, bone marrow was flushed in PBS from the femur and tibia of mice. Cells were cultured in bacterial grade plastic for 7 days at 37°C, 5% CO_2_. Cells were then detached using Versene (Gibco), re-plated on tissue culture plastic in the appropriate media and used within 48h. Primary cells were cultured in DMEM supplemented with 10% heat inactivated FBS, 2 mM L-glutamine, 100 U/ml penicillin G, 100 μg/ml streptomycin and 0.25 μg/ml amphotericin. BMDMs were cultured in 5 ng/ml M-CSF while GM-BMCs were cultured in 10 ng/ml GM-CSF. Where indicated cells were treated with 2 μM MRT67307, 2 μM PD184352, 1 μM VX-745 of 10 μM SB-74651A for 1 h before stimulation. Cells were stimulated with either 100 ng/ml LPS (*E. coli 055:B5*) or 10 µM PGE2 for the times indicated in the legends.

For CRTC knockdown experiments, siRNA was transfected into primary BMDMs using the Lonza Amaxa Optimized Mouse Macrophage Nucleofection system (catalog number VPA-1009). Smartpool siGENOME siRNAs were purchased from Thermo-Dharmacon. These experiments gave efficient knockdown of CRTC1, 2 and 3 as reported previously [29].

### qPCR

Following stimulation cells were lysed and total RNA isolated using Nucleospin RNA purification columns according to the manufactures protocol (Macherey-Nagel). 0.5 to 1 µg of total RNA was reverse transcribed using BioRad iScript, qPCR was then carried out using SyberGreen based detection methods (Takara Biosciences). Fold induction relative to the wild type unstimulated sample was calculated as described using 18s and/or GAPDH as reference gene [36]. Primer sequences for 18s, GAPDH, CRTCs, IL-12p40 and IL-12p70 have been described previously [29, 72]. IL-23p19 mRNA was detected using the primers ATCCAGTGTGAAGATGGTTGTGAC and TTCTAGTAGGGAGGTGTGAAGTTG while IL-23p19 primary transcript was detected with the primers AATGTGCCCCGTATCCAGTG and R: GGCTTAGTGGTACCTGGCTG. The primers for IL-17A were CTCCAGAAGGCCCTCAGACTAC and AGCTTTCCCTCCGCATTGACACAG and IL-17F were CTGGAGGATAACACTGTGAGAGT and TGCTGAATGGCGACGGAGTTC.

### Cytokine measurements

Cytokines were measured from serum or cell culture media as described. For measurement of cytokine levels in inflamed skin, tissue from an ear notch (2 mm diameter) was homogenised in protein lysis buffer (50mM Tris HCl (pH 8), 150mM NaCl, 0.5% Sodium Deoxycholate, 1%NP-40, 0.1% SDS, 1µg/ml Leupetin, 1µg/ml Aprotinin, 1mM PMSF) using 5mm Stainless Beads and TissueLyser II (frequency: 30/sec, duration 2 × 30 sec) (Qiagen, Hilden, Germany). IL-23 was measured using either a mouse IL-23 ELISA (R&D) or Luminex based method from BioRad. IL-12p40, Il-12p70, IL-17A, IL-17F and IL-10 were measured using a Luminex based method using Bioplex reagents from BioRad.

### Immunoblotting

Cells were lysed directly into SDS sample buffer and aliquots run on 10% polyacrylamide gels using standard methods. Proteins were transferred onto nitrocellulose membranes, and specific proteins were detected by immunoblotting. Antibodies against phospho-ERK1/2, phospho-p38, total ERK1/2, total p38α, phospho-Ser133 CREB (which also recognizes ATF1 phosphorylated on Ser63) and phospho-Thr581 MSK1 were from Cell Signalling Technology. The total MSK1 antibody was generated in-house [22]. HRP-conjugated secondary antibodies were from Pierce (Cheshire, U.K.), and detection was performed using the ECL reagent from Amersham Biosciences (Buckinghamshire, U.K.).

### Imiquimod induced skin inflammation

Mice were treated with Aldara cream (5% Imiquimod, Meda SD, Solna, Sweden) or Oilatum (control soft cream, Glaxo Operation UK Ltd, Durham, UK). All experiments were carried out blind to the genotype of the animals. Aldara and Oilatum were applied for 7 consecutive days; equally on dorsal and ventral surface of right and left ears with a total dose of 40mg total per mouse per day. Animals were maintained on a 12h light / 12 h dark cycle, and cream was applied between 3 and 4 h into the light cycle. Once daily, prior to application, ear thickness was measured using a Mitutoyo 7301 dial thickness gauge (Mitutoyo, Kawasaki, Japan). Mice were sacrificed 4 hours or 3 days after last treatment as indicated in the figure legends, and ear samples were taken for further analysis. If required, an intermediate 2mm diameter ear biopsy was taken during the experiment. Samples for mRNA expression level and protein analysis were snap frozen in liquid nitrogen and stored at −80”C until further use.

### Flow cytometry

Spleens were isolated from mice and single cell suspensions prepared via passing through a 100 μm filter. After RBC lysis cells were stained with 0.5 μg/ml DAPI and live cells were counted on BD FACSVerse ™. For flow cytometry analysis, 1×10^6^ cells were blocked for 20 min at 4°C with FcR antibody (purified anti-CD16/32; BD Pharmingen) diluted (1:50) in PBS containing 1% Bovine serum albumin. For neutrophils and monocytes characterisation, the cells were stained for 20 min at 4°C with the anti-CD45 (BV510), anti-NK1.1 (APC/Cy7), anti-CD11b (PE/Cy7), anti-Gr-1 (PerCp/Cy5.5), anti-CD115 (APC), anti-Ly-6C, anti-CX3CR1(PE), anti-I-A/I-E (MHCII) (AlexaFluor700), anti-CD11c (PE/Dazzle)/ To measure intracellular levels of IL-17, cells were seeded at density of 1×10^6^ in 1mL complete RPMI media (RPMI-1640 medium with 10% heat-inactivated FBS, 50 U/ml penicillin-streptomycin, 10mM HEPES buffer, 1mM sodium pyruvate, 50 µM 2-mercaptoethanol, 2mM L-Glutamine, 0.1mM Non-essential amino acids) and stimulated with 50ng/ml PMA and 1µg/ml ionomycin plus brefeldin A (5 μg/ml)for 4 hours.

Following stimulation cells were washed with FACS buffer (PBS + 1% BSA), fixed using fixation buffer (eBioscience) for 20min at +4°C, washed with FACS buffer, permeabilised with 1x permeabilisation buffer (eBioscience) for 20min at +4°C, After washing in FACS buffer cells were incubated with 1:50 Fc block made up in 1x permeabilisation buffer for 20min at +4°C, washed with FACS buffer and incubated with antibodies (1:100 TCR-β FITC, 1:100 TCR-γδ APC, 1:200 IL-17 PE, and 1:200 CD4 PerCp/Cy5.5) made up in 1x permeabilisation buffer for 30min at +4°C. Stained cells were washed with FACS buffer and data acquired on a BD LSR Fortessa™ and further analysed using FlowJo software.

## Author contributions

KFH, MGB, TP and SN carried out experimental work, all authors contributed to data analysis and preparation of the manuscript.

## Funding and additional information

The work was supported by grants from the Medical Research Council and Versus Arthritis (JSCA)

## Conflict of interest

The authors have no conflict of interest.

## Notes

### Competing Interest Statement

The authors have declared no competing interest.

