## supplemental data for "IL-23 production is regulated via an MSK1/2 – CREB dependent signalling pathway downstream of Toll like receptors"

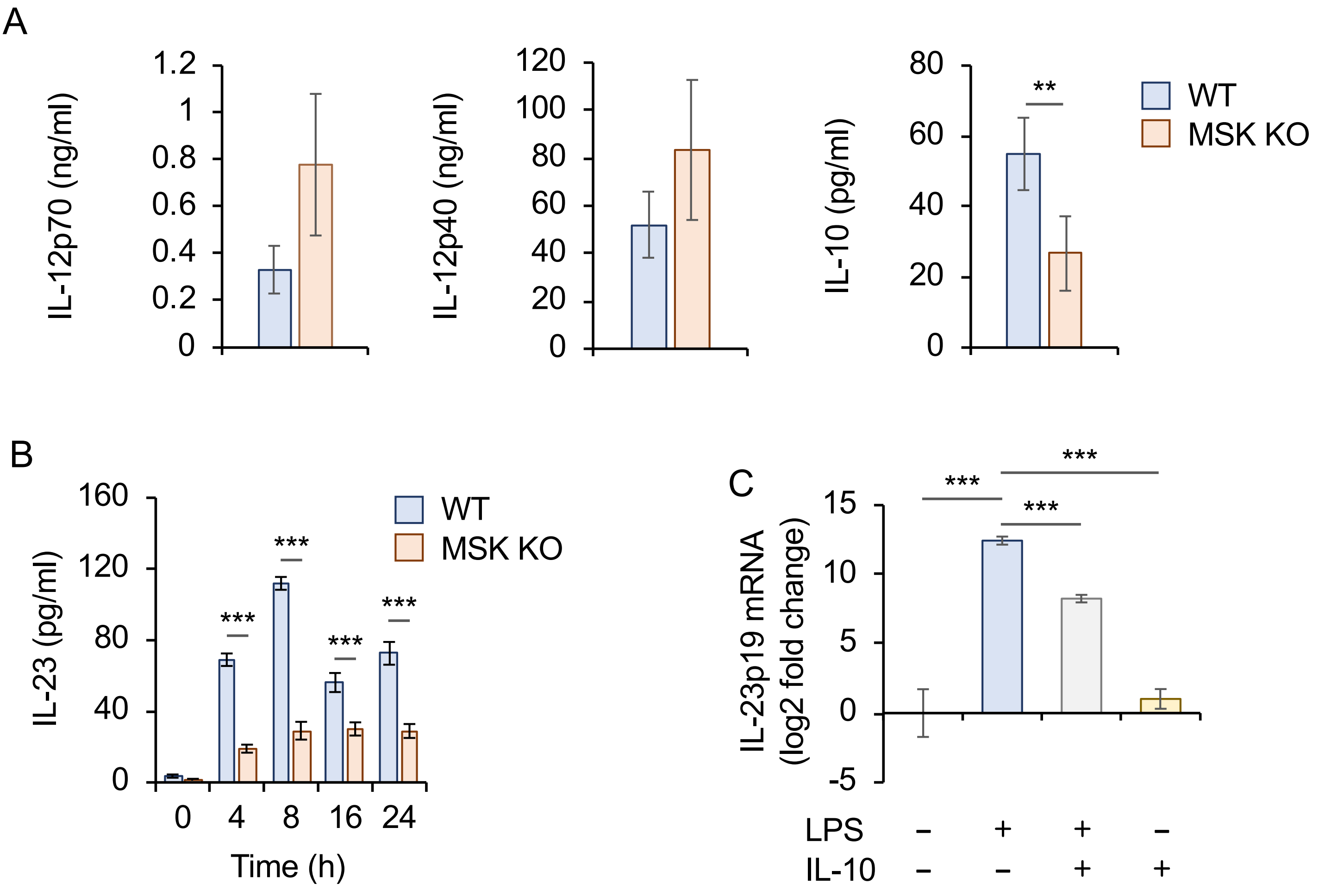


**Supplementary Figure 1**

A) Wild type or MSK1/2 knockout GM-BMCs were stimulated for the 8h with 100 ng/ml LPS and the levels of IL-12p70, IL-14p40 and IL-10 secreted into the media were determined by a Luminex based assay.

B) Wild type or MSK1/2 knockout BMDMs were treated for 24 h with 5ng/ml GM-CSF and then stimulated for the indicated times with 100 ng/ml LPS. IL-23 secretions was determined by ELISA.

C) Wild type BMDMs were stimulated with 100ng/ml LPS and or 10ng/ml IL-10 for 1 h as indicated. IL-23p19 mRNA was determined by qPCR.

For all panels, n=4, with $p$<0.01 indicated by ** and $p$<0.001 by ***. For A and C a two tailed unpaired Students ttest was used while a 2 way ANOVA with Holm-Sidak post hoc testing was used for B.

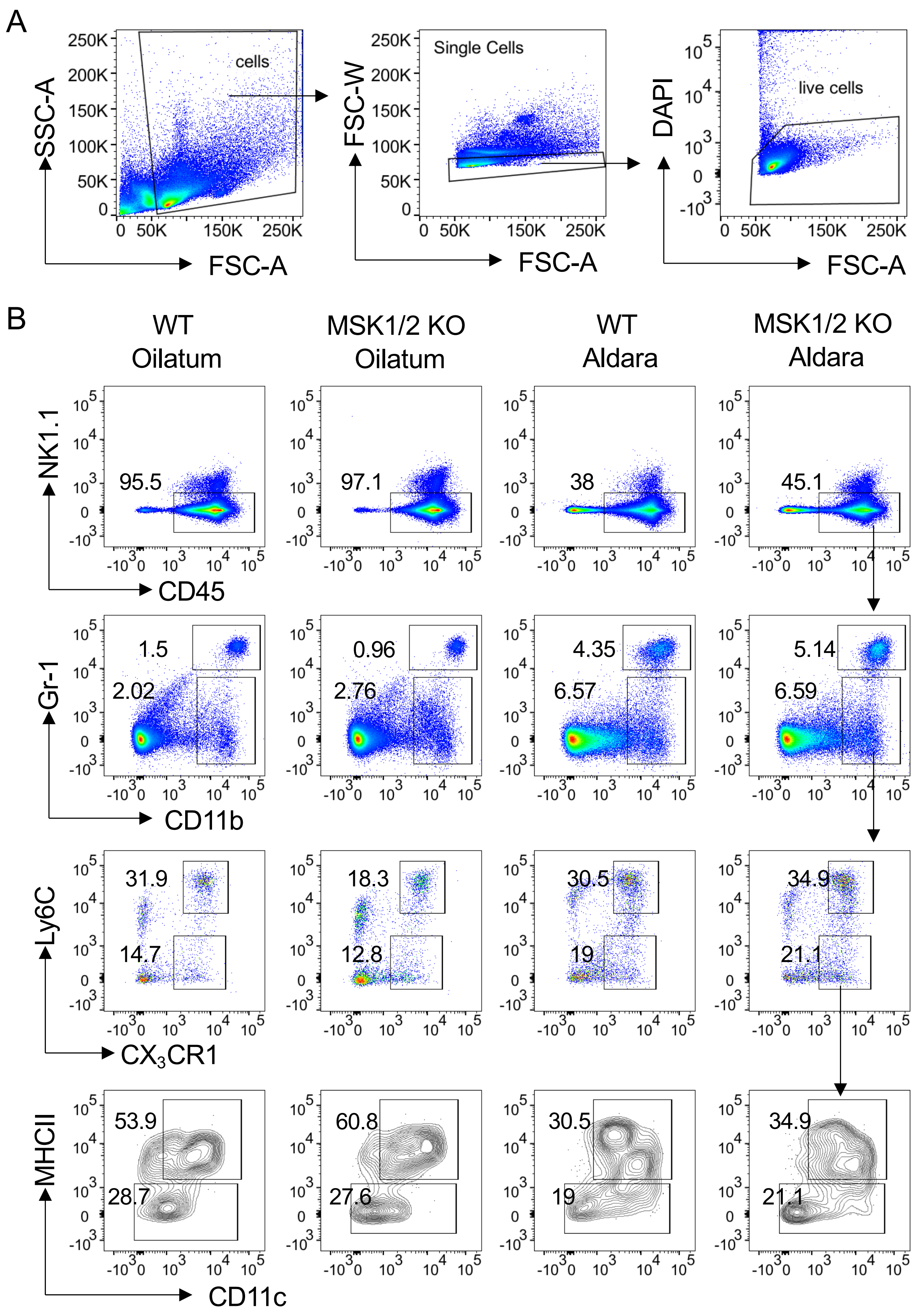


**Supplementary figure 2**

Wild type or MSK1/2 knockout mice were treated daily with Aldara or Oilatum cream for 6 days. Spleens were isolated and analysed by flow cytometry as described in the methods. (A) Cells were gated based on FSC-A and SSC-A, doublets were excluded based of FSC-A and FSC-W and DAPI staining was used to gate out dead cells. (B) Representative flow cytometry plots are showing gating strategy for identifying neutrophils (CD45$^{+ve}$NK1.1$^{-ve}$Gr-1$^{high}$), Ly6C$^{+ve}$ inflammatory (CD45$^{+ve}$NK1.1$^{-ve}$Gr-1$^{low/-ve}$CX3CR1$^{+ve}$Ly6C$^{+ve}$) and Ly6C$^{-ve}$ patrolling monocytes (CD45$^{+ve}$NK1.1$^{-ve}$Gr-1$^{low/-ve}$CX3CR1$^{+ve}$Ly6C$^{-ve}$MHCII$^{-ve}$)

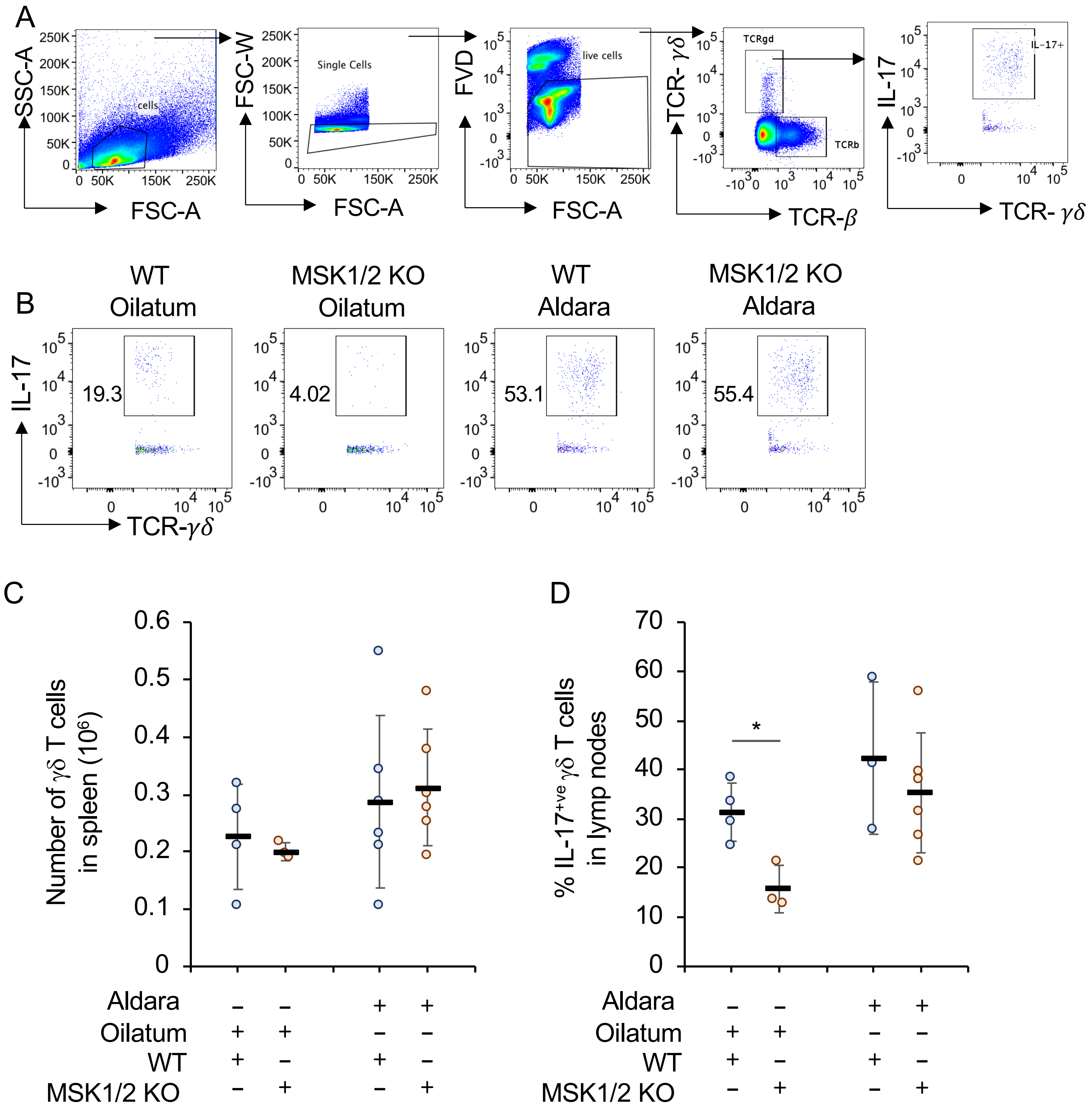


**Supplementary figure 3**

A-B) Wild type or MSK1/2 knockout mice were treated daily with Aldara or Oilatum cream for 6 days. Spleens were isolated and cells restimulated with PMA and ionomycin before staining to identify IL-17$^{+ve}$ γδ T cells as described in the methods. (A) Gating strategy and representative flow cytometry plots for the identification of IL-17$^{+ve}$ γδ T cells. (B) Total γδ T cells in the spleens of wild type and MSK1/2 knockout mice. C) Lymph nodes were isolated from mice that were treated daily with Aldara or Oilatum cream for 6 days. Cells were stimulated with PMA and ionomycin and the percentage of IL-17$^{+ve}$ γδ T cells determined. Plots show the average and standard deviation in each of the experimental groups with individual mice shown by circles. $p$<0.05 (two tailed unpaired Students ttest) is indicated by *.
